# Image-Based Assessment of Natural Killer Cell Activity against Glioblastoma Stem Cells

**DOI:** 10.1101/2023.12.29.573629

**Authors:** Yuanning Du, Samuel Metcalfe, Shreya Akunapuram, Angela M. Richardson, Aaron A. Cohen-Gadol, Jia Shen

**Author notes:** Correspondence: J Shen, Medical Sciences Program, Indiana University School of Medicine, 202 Biology Building, 1001 E. 3rd St., Bloomington, IN 47405, USA.

## Abstract

Glioblastoma (GBM) poses a significant challenge in oncology and stands as the most aggressive form of brain cancer. A primary contributor to its relentless nature is the stem-like cancer cells, called glioblastoma stem cells (GSCs). GSCs have the capacity for self-renewal and tumorigenesis, leading to frequent GBM recurrences and complicating treatment modalities. While natural killer (NK) cells exhibit potential in targeting and eliminating stem-like cancer cells, their efficacy within the GBM microenvironment is limited due to constrained infiltration and function. To address this limitation, novel investigations focusing on boosting NK cell activity against GSCs are imperative. This study presents two streamlined image-based assays assessing NK cell migration and cytotoxicity towards GSCs. It details protocols and explores the strengths and limitations of these methods. These assays could aid in identifying novel targets to enhance NK cell activity towards GSCs, facilitating the development of NK cell-based immunotherapy for improved GBM treatment.

## Introduction

Glioblastoma (GBM), the most frequent primary malignant brain tumor in adults, is known for its aggressiveness and lethality. The standard treatment protocol for GBM involves a combination of surgical intervention to remove as much of the tumor as feasible, followed by a course of radiotherapy and chemotherapy [1]. However, due to the recurrent nature of GBM, these treatments exhibit limited efficacy, resulting in notably low survival rates - reportedly less than 10% over a five-year period in the United States [1]. The resistance of GBM to standard treatments primarily arises from intratumoral heterogeneity, principally driven by glioblastoma stem cells (GSCs), which constitute a population of stem-like cancer cells within the tumors [2]. These GSCs, possessing properties akin to normal stem cells such as self-renewal and differentiation, play a pivotal role in GBM initiation, resistance to therapy, and the aggressive relapse of this form of brain cancer [3]. Unless the GSCs are eradicated, achieving a cure for GBM remains improbable.

The human immune system is divided into the innate and adaptive immunity, working together to protect against infections and cancers. The innate system, including natural killer (NK) cells and neutrophils, serves as the body’s first line of defense, swiftly responding to potential threats [4-7]. This system works alongside the adaptive immune system, including T cells and B cells, providing prolonged immunity through antibody responses and cell-mediated immune responses. NK cells within the innate system can be activated via diverse receptors, including MHC-I related (e.g., MICA or MICB) and non-related (e.g., Nectin-2 or CD155) receptors, enabling swift recognition and elimination of virus-infected or tumor cells [8]. Previous studies demonstrate the capability of NK cells to recognize and destroy stem-like cancer cells in various cancer types in vitro [9, 10]. However, in GBM, the immunosuppressive tumor microenvironment restricts NK cell function, impacting their infiltration and cytotoxicity [11-13]. Further exploration is needed to investigate the promising therapeutic potential of enhancing NK cell activity against GSCs within this environment for eradicating GSCs and treating GBM [14].

Based on this background, our study aims to develop two efficient image-based assays evaluating NK cell migration and cytotoxicity towards GSCs. These assays provide sensitive, direct, and reproducible methods and could contribute to identifying new targets for boosting NK cell efficacy against GSCs. This could advance the development of NK cell-based immunotherapy for enhanced GBM treatment.

### Materials

1. NK-92MI (ATCC, Cat # CRL-2408)
2. MyeloCult H5100 culture media (STEMCELL Technologies, Cat # 05150)
3. Hydrocortisone (STEMCELL Technologies, Cat # 74142)
4. GSCs (a gift from Dr. Jeremy Rich at the UPMC Hillman Cancer Center, Pittsburgh, PA, USA)
5. Neurobasal media (Gibco, Cat # 12349015)
6. B27 without vitamin A (Gibco, Cat # 12587010)
7. EGF (R&D Systems, Cat # 236-EG-01M)
8. bFGF (R&D Systems, Cat # 3718-FB-025)
9. Sodium pyruvate (Gibco, Cat # 11360070)
10. Glutamax (Gibco, Cat # 35050061)
11. Penicillin/streptomycin (Cytiva, Cat # SV30010)
12. Lenti-X-293T cells (TaKaRa Bio, Cat # 632180)
13. DMEM media (Gibco, Cat # 11995065)
14. FBS (Corning, Cat # 35015CV)
15. Accutase (Innovative Cell Technologies, Cat # AT104500)
16. DPBS, no calcium, no magnesium (Gibco, Cat #14190250)
17. DMSO (Fisher BioReagents, Cat # BP2311)
18. Control and gene-X shRNAs vectors (gifts from Dr. Charles Spruck, Sanford Burnham Prebys Medical Discovery Institute, La Jolla, CA, USA)
19. jetPRIME transfection reagent (Polyplus, Cat # 101000001)
20. Transwell inserts with diameter 6.5 mm, pore size 5 mm (Corning, Cat # 3421)
21. Calcein AM (eBioscience, Cat # 65085339)
22. U-Shaped-Bottom 96-well plate (Fisherbrand, Cat # FB012932)
23. Hemocytometer (Weber Scientific, Cat # 304811)
24. Axiovert 40 CFL Inverted Microscope (Zeiss)
25. X-Cite 120 Fluorescence Illumination Systems (EXFO Photonic Solutions Inc.)

### Methods

#### Cell culture

Prepare the NK-92MI cell culture media by adding 500 uL of 10 mM hydrocortisone and 5 mL of penicillin/streptomycin into 500 mL of MyeloCult H5100 culture media. Maintain the NK-92MI cells expressing human IL-2 in NK cell culture media in a 5% CO2, 37 °C incubator, passaging them every 2 - 3 days to retain a cell density between 2 × 10^5^ and 8 × 10^5^ cells/ml. For Lenti-X-293T cells, utilize DMEM media supplemented with 10 % FBS for cell culture. Regarding GSCs culture, employ Neurobasal media supplemented with B27 lacking vitamin A, 20 ng/ml EGF, 20 ng/ml bFGF, 1% sodium pyruvate, 1% GlutaMAX, and 1% penicillin/streptomycin [15].

#### Lentivirus preparation, infection, and GSC conditioned media collection

The lentivirus packaging plasmids (pMD2.g and psPAX2) alongside control or gene-X shRNA vectors are transfected into Lenti-X-293T cells cultured in DMEM media using jetPRIME transfection reagent. Following 24 h, the DMEM media is replaced with GSC culture media and incubated at 37 °C for an additional 48 h. The collected supernatant containing lentivirus undergoes filtration through 0.45 μm filters before GSC infection. To infect GSCs, incubate the lentivirus expressing either control or gene-X shRNA with GSCs for 24 h, followed by a media change for another 48 h. Gather the conditioned media from these GSCs for NK cell migration assay. Simultaneously, portion the surplus conditioned media into 1.5 mL tubes and store them at -80 °C for future utilization. The GSCs transduced with either control or gene-X shRNA are collected for GSC-NK co-culture and NK cell cytotoxicity assay.

#### Image-based NK cell migration assay

1. Precondition transwell inserts by introducing 600 µL of freshly prepared GSC media into the lower compartment of 24-well plate wells, followed by the placement of inserts. Subsequently, add 100 µL of NK cell media into the upper compartment of the inserts. Incubate the setup for 1 h in a 5% CO2, 37 °C incubator.
2. Harvest NK-92MI cells into a 15 mL centrifuge tube, subjecting them to centrifugation at 500 x g for 5 min at 4 °C.
3. Aspirate the media and resuspend the cells in 1 mL of fresh NK cell media. Subsequently, quantify the NK-92MI cells using a hemocytometer.
4. Retrieve the preconditioned 24-well plate from the incubator. Remove 600 µL of media from the lower compartment of each well and replace it with 600 µL of GSC conditioned media collected when culturing GSCs transduced with control or gene-X shRNA, as detailed in the “Lentivirus preparation, infection, and GSC conditioned media collection” section.
5. Place the preconditioned transwell inserts into these wells and withdraw the initial 100 µL of NK cell media. Add 100 uL of 10^5^ NK-92MI cells into each transwell insert.
6. Include fresh GSC media in lower compartment of 24-well plate wells with NK-92MI cells in the inserts, serving as negative controls.
7. Incubate the plate for 4 h in a 5% CO2, 37 °C incubator.
8. Extract the inserts and examine the lower compartment of the wells using a Zeiss Axiovert 40 CFL inverted microscope. Any migrated NK-92MI cells should be observable in the wells.
9. Capture images of cells in the wells using ZEN blue software connected to the microscope at 4 x magnification and quantify the migrated NK-92MI cells in different fields.

Please refer to the schematic of this assay in Fig.1.

**Fig 1.**
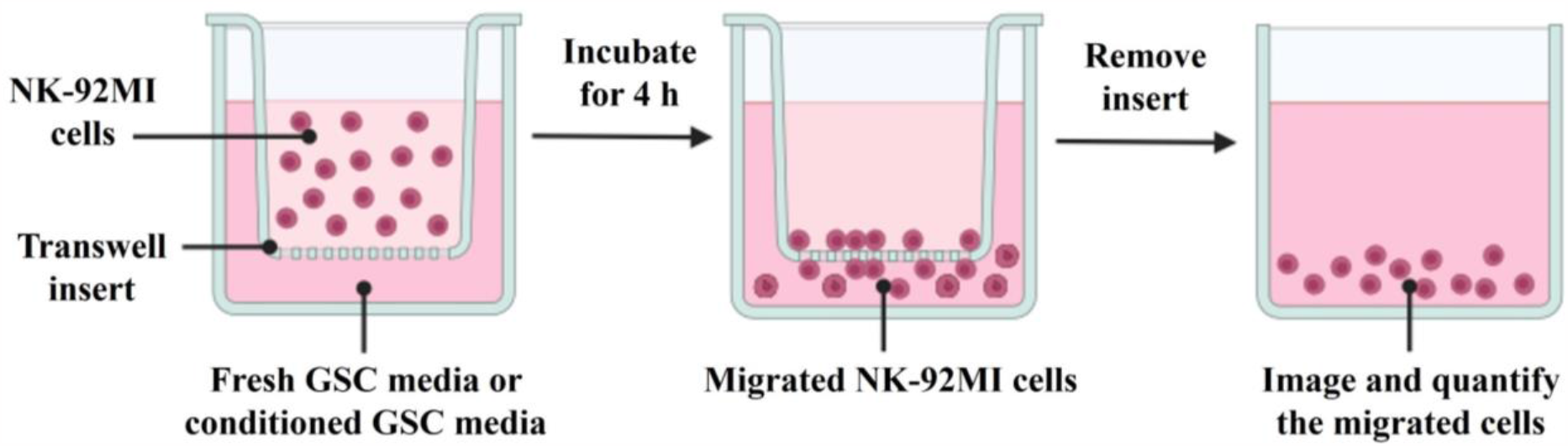
Schematic of the image-based NK cell migration assay.

#### Image-based NK cell cytotoxicity assay

1. Centrifuge GSCs previously transduced with either control or gene-X shRNA as detailed in the “Lentivirus preparation, infection, and GSC conditioned media collection” section at 500 x g for 5 min.
2. Resuspend the resulting cell pellets in 1 mL of 0.25% accutase and incubate at 37 °C for 5 min to obtain a single-cell suspension.
3. Subject the cell suspension to another round of centrifugation at 500 x g for 5 min. Resuspend the cells in 3 mL of GSC culture media.
4. Add 1.5 µL of calcein AM solution (10 mM stock in DMSO) to the cells and gently mix. Incubate the mixture for 30 min in a 5% CO2, 37 °C incubator.
5. Centrifuge the calcein AM-labelled GSCs at 500 x g for 5 min.
6. Wash the cells 3 times with 5 mL of DPBS to remove excess calcein AM dye.
7. Centrifuge NK-92MI cells at 500 x g for 5 min. Resuspend them in fresh NK cell media.
8. Utilize a hemocytometer to count both the calcein AM-labelled GSCs and the NK-92MI cells.
9. Adjust the GSC concentration to 10^5^ cells/mL in GSC culture media and the NK-92MI cells to 2 x 10^6^ cells/mL in NK cell media.
10. In a U-Shaped-Bottom 96-well plate, place calcein AM-labelled GSCs (10^4^ cells/100 µL/well) with or without NK-92MI cells (2 x 10^5^ cells/100 µL/well) in duplicate. Utilize GSCs (control vs. gene-X knockdown) only with 100 µL NK cell culture media as controls.
11. Incubate the cells for 4 h in a 5% CO2, 37 °C incubator.
12. Post-incubation, capture fluorescence images for the calcein AM signal (green) in the wells using a fluorescence microscope (Axiovert 40 CFL Inverted Microscope) connected to X-Cite 120 Fluorescence Illumination Systems at 10 x magnification.
13. Quantify the calcein AM signal in the images. Calculate the percentage viability of GSCs using the formula: Viability % = (# of “green” GSCs co-cultured with NK-92MI cells / # of “green” GSCs in controls) x 100 Please refer to the schematic of this assay in Fig.2.

**Fig 2.**
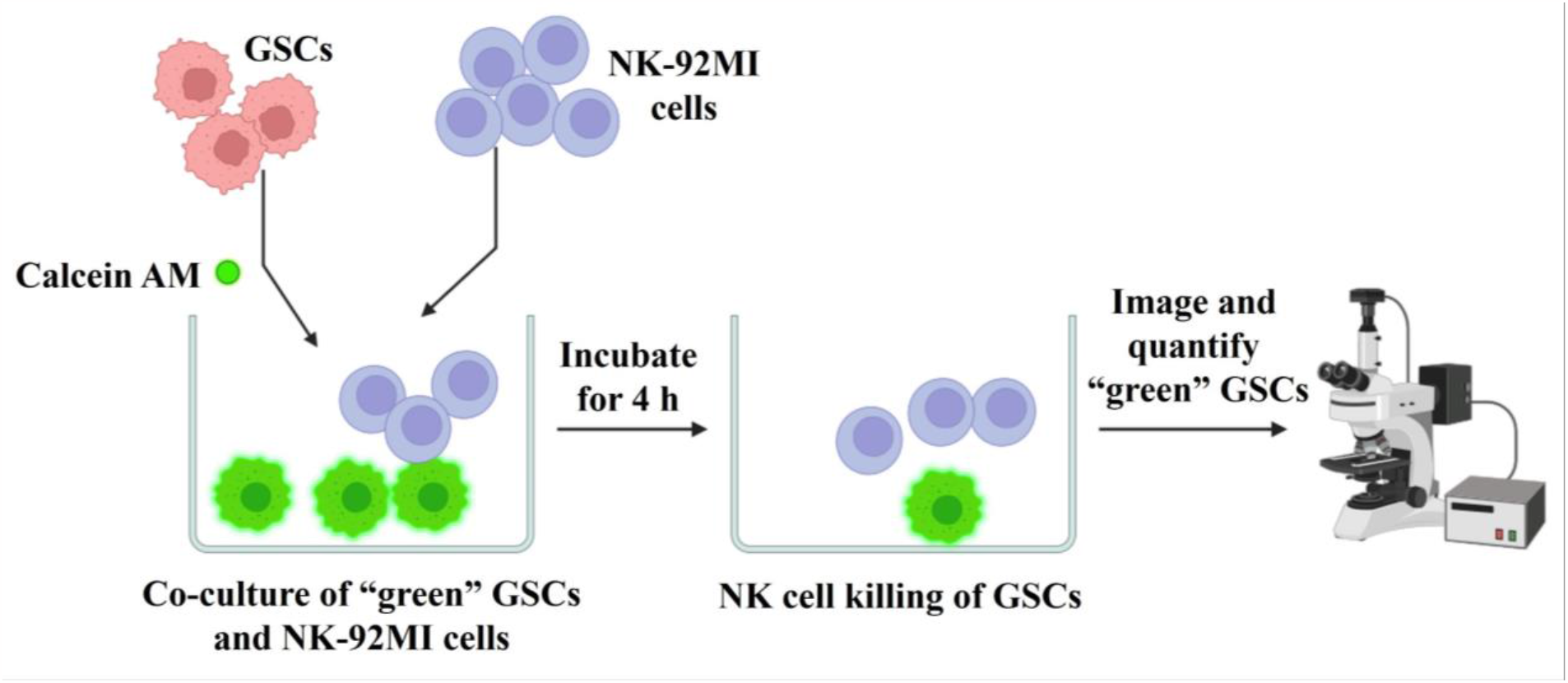
Schematic of the image-based NK cell cytotoxicity assay.

## Results and discussion

In the NK cell migration assay, three distinct groups were examined: a control group with fresh GSC culture media, a conditioned media group from GSCs infected by lentivirus expressing control shRNA, and an experimental group with conditioned media from GSCs exhibiting gene-X knockdown (Fig. 3A). Quantification of migrated NK-92MI cells was performed by analyzing images acquired from the lower compartment of the wells. The results indicate a significant increase in the migration of NK-92MI cells towards the conditioned media from GSCs with gene-X knockdown (Fig. 3B). Indeed, our previous findings suggested that the depletion of gene-X in GSCs could provoke heightened expression of various cytokines responsible for attracting NK cells, consequently stimulating NK cell migration towards the GSCs. This affirms the credibility and validity of our assay methodology.

**Fig 3.**
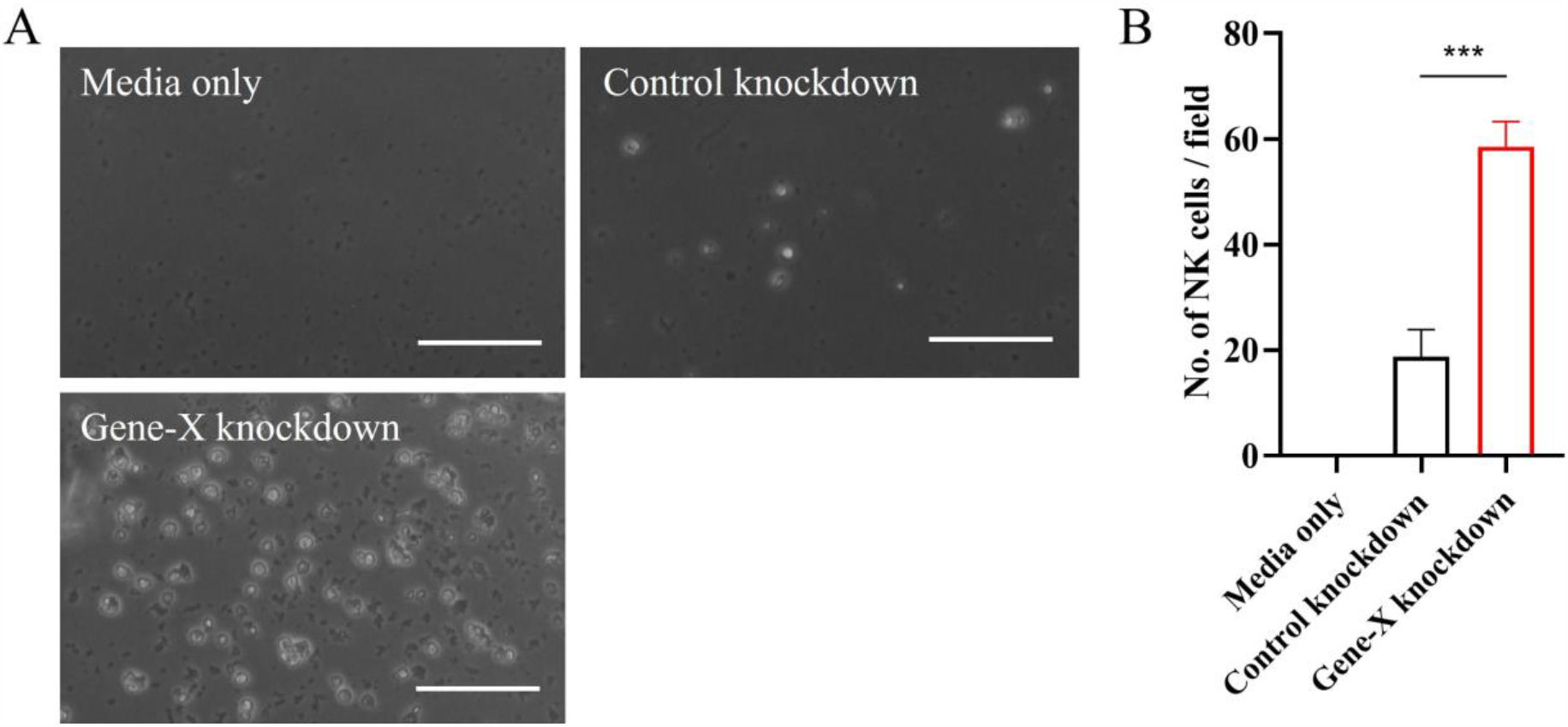
(A) Representative images of the NK-92MI cells that migrated into the lower compartment of the 24-well plate. Scale bar, 100 μm. (B) Quantification of the migrated NK-92MI cells in different fields (n = 4). Data represent mean ± SEM. ***p < 0.001 by one-way ANOVA followed by Tukey’s multiple comparisons test.

In the NK cell cytotoxicity assay, GSCs subjected to control conditions or gene-X knockdown were initially labelled using calcein-AM before their co-culture with or without NK-92MI cells (Fig. 4A). The results reveal a marginal decline in the viability of GSCs following co-culture with NK-92MI cells, suggesting limited cytotoxicity exerted by the NK cells against GSCs. However, the knockdown of gene-X in the GSCs notably augmented the cytotoxic effects of NK-92MI cells on the GSCs (Fig. 4B). Correspondingly, our initial observations suggested that the knockdown of gene-X could elevate the expression of various NK cell activating receptors on GSCs, thereby triggering the recognition and subsequent elimination of GSCs by NK cells. Consequently, these findings further validate the reliability and precision of our assay method.

**Fig 4.**
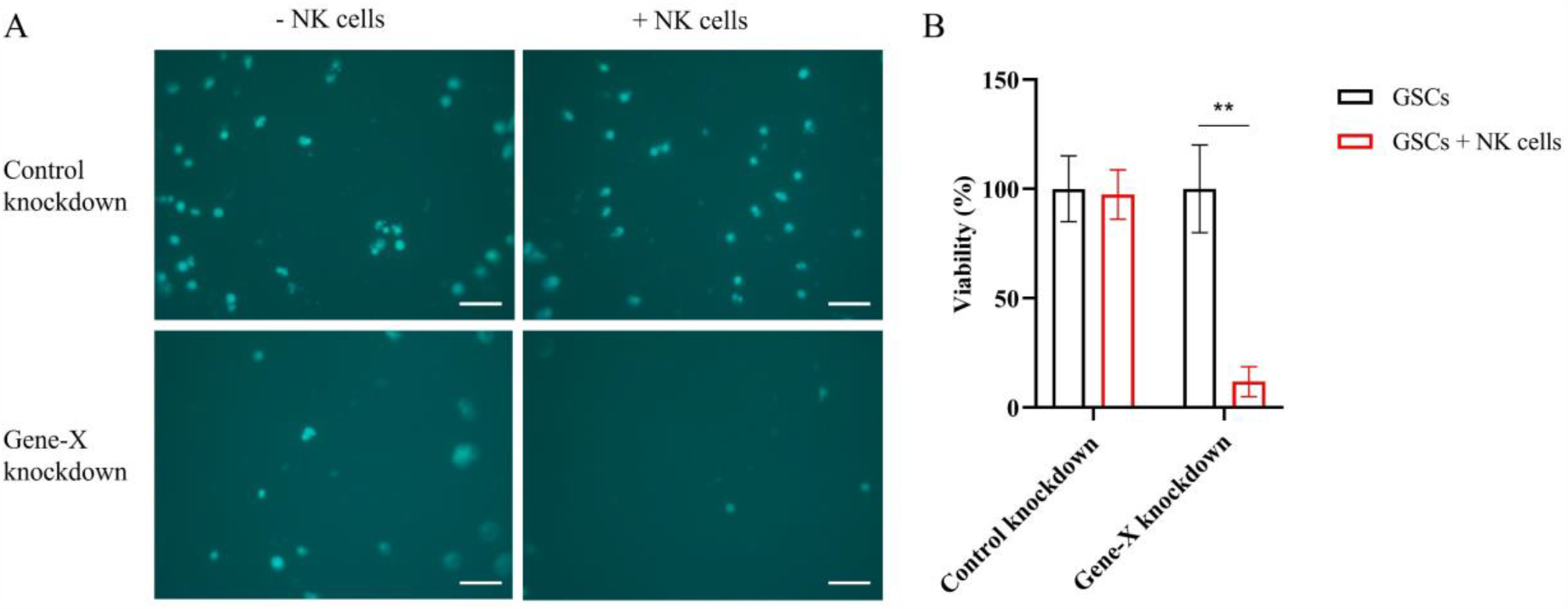
(A) Representative fluorescent images of calcein AM-labelled GSCs. Scale bar, 100 μm. (B) Quantification of the viability of GSCs in different fields (n = 4). Data represent mean ± SEM. **p < 0.01 by two-way ANOVA followed by Sidak’s multiple comparisons test.

Patient-derived GSCs at an early passage were employed in the assays, underscoring the clinical significance and relevance of this research. This article delineates two image-based methodologies aimed at efficiently examining NK cell migration and cytotoxicity towards GSCs. To quantify the migrated NK cells, unlike traditional approaches such as flow cytometry, which require expensive equipment and specialized training [16-18], our detection method is straightforward, cost-effective, and adaptable to basic microscopy available in most laboratory setups, reducing operational complexities. Traditionally, the chromium release assay has been a mainstay for evaluating NK cell cytotoxicity [16, 19]. However, its reliance on radioisotopes like chromium-51 presents health risks, demands specific training, and involves handling expensive and potentially hazardous radioactive materials. Furthermore, the assay’s dependency on spontaneous release may yield inconsistent results. In contrast, our outlined protocol offers a simplified, easily implementable alternative applicable to laboratories equipped with a fluorescent microscope. Moreover, both of our assays enable direct quantification of cells (NK-92MI cells in the migration assay and GSCs in the cytotoxicity assay) within the experimental wells, obviating the need for transfer or additional instrumentation. This not only saves time but also minimizes potential variations introduced during such additional processing steps. Additionally, it’s important to acknowledge the limitations here. While aiding in comprehending NK cell behavior towards GSCs, these assays may not entirely mirror the complexity of in vivo settings. For instance, the influence of other immune cell players, such as T cells or macrophages, on the effects exerted by NK cells against GSCs in GBM microenvironment remains a pertinent question. Consequently, validating the outcomes obtained in vitro through subsequent in vivo investigations is imperative. This strategic approach would bolster the practical applicability of the insights garnered from these assays.

Collectively, this research introduces methodologies that are straightforward, dependable, and adaptable for application in diverse laboratory environments, enabling the investigation of NK cell migration and cytotoxicity against GSCs. While acknowledging the study’s limitations, the outlined methodology holds promise in delineating the feasibility of identifying novel targets and leveraging NK cell immunotherapy for targeting GSCs and treating GBM.

### Tips & Tricks

1. Establishing an initial equilibration phase for the transwell inserts is crucial for preserving the optimal functionality of NK cells in the migration assay. This process entails a stepwise addition of GSC media to the wells within a 24-well plate, followed by the insertion of transwell inserts, and subsequent addition of NK cell media into the inserts. A recommended minimum incubation period of 1 h for the plate is advised to achieve the desired equilibrium.
2. The inserts are supplied in packs of 12, prearranged within a 24-well plate. If fewer than 12 migrations are being conducted, it’s possible to extract the required number of inserts, utilizing a standard 24-well plate, and then reseal the remaining inserts in their container.
3. Employing U-Shaped-Bottom 96-well plates for the cytotoxicity assay might offer an improved spatial proximity conducive to the interaction between NK-92MI cells and GSCs compared to regular flat-bottom plates.
4. To optimize calcein AM staining of GSCs, conducting a preliminary experiment that varies concentrations and incubation times is advisable.
5. The use of DPBS for washing GSCs subsequent to calcein AM staining is crucial to minimize nonspecific high background staining.
6. Given the distinct culture media requirements for NK-92MI cells and GSCs, extended co-culture periods have the potential to affect their cellular traits and behaviors during analyses. Therefore, it is advisable to minimize prolonged co-culture durations to uphold the integrity of experiments and mitigate the likelihood of changes in cellular characteristics.

## Acknowledgements

Research reported in this publication was supported by the National Cancer Institute of the National Institutes of Health under Award Number P30CA030199. The content is solely the responsibility of the authors and does not necessarily represent the official views of the National Institutes of Health. The authors express their gratitude to the following for their valuable contributions: the Genomics Core Facility and Histology Core Facility, and Dr. Charles Spruck’s lab at the Sanford Burnham Prebys Medical Discovery Institute for their research support. Additionally, appreciation is extended to Dr. Jeremy Rich at the UPMC Hillman Cancer Center for providing the GSCs, and Dr. Kenneth Nephew, Dr. Claire Walczak, and Dr. Peter Hollenhorst at the Indiana University School of Medicine for their assistance in optimizing the image-based assays. Fig.1 and Fig.2 were created with BioRender.com.

## Abbreviations

GBM: glioblastoma
GSCs: glioblastoma stem cells
NK cells: natural killer cells
MHC-I: major histocompatibility class I
EGF: epidermal growth factor
bFGF: basic fibroblast growth factor
IL-2: interleukin-2
FBS: fetal bovine serum
DPBS: Dulbecco’s Phosphate-Buffered Saline
DMEM: Dulbecco’s Modified Eagle Medium
DMSO: dimethyl sulfoxide
ATCC: American Type Culture Collection

## Author contributions

YD, SM, and JS performed the experiments. YD, SM, SA, AMR, AACG and JS contributed to writing, reviewing, and editing the manuscript. JS conceived and designed the project.

## Data accessibility

The data included in the figures serve solely for illustrative purposes and are part of an ongoing study that is in the process of publication.

## References

1. Schaff, L. R. & Mellinghoff, I. K. (2023) Glioblastoma and Other Primary Brain Malignancies in Adults: A Review, Jama. 329, 574–587.

2. Prager, B. C., Bhargava, S., Mahadev, V., Hubert, C. G. & Rich, J. N. (2020) Glioblastoma Stem Cells: Driving Resilience through Chaos, Trends in cancer. 6, 223–235.

3. Gimple, R. C., Bhargava, S., Dixit, D. & Rich, J. N. (2019) Glioblastoma stem cells: lessons from the tumor hierarchy in a lethal cancer, Genes & development. 33, 591–609.

4. Kraus, R. F. & Gruber, M. A. (2021) Neutrophils-From Bone Marrow to First-Line Defense of the Innate Immune System, Frontiers in immunology. 12, 767175.

5. Nicholson, L. B. (2016) The immune system, Essays in biochemistry. 60, 275–301.

6. Yatim, K. M. & Lakkis, F. G. (2015) A brief journey through the immune system, Clinical journal of the American Society of Nephrology : CJASN. 10, 1274–81.

7. Lupo, K. B. & Matosevic, S. (2019) Natural Killer Cells as Allogeneic Effectors in Adoptive Cancer Immunotherapy, Cancers. 11.

8. Wu, S. Y., Fu, T., Jiang, Y. Z. & Shao, Z. M. (2020) Natural killer cells in cancer biology and therapy, Molecular cancer. 19, 120.

9. Ames, E., Canter, R. J., Grossenbacher, S. K., Mac, S., Chen, M., Smith, R. C., Hagino, T., Perez-Cunningham, J., Sckisel, G. D., Urayama, S., Monjazeb, A. M., Fragoso, R. C., Sayers, T. J. & Murphy, W. J. (2015) NK Cells Preferentially Target Tumor Cells with a Cancer Stem Cell Phenotype, J Immunol. 195, 4010–9.

10. Luna, J. I., Grossenbacher, S. K., Murphy, W. J. & Canter, R. J. (2017) Targeting Cancer Stem Cells with Natural Killer Cell Immunotherapy, Expert opinion on biological therapy. 17, 313–324.

11. Balatsoukas, A., Rossignoli, F. & Shah, K. (2022) NK cells in the brain: implications for brain tumor development and therapy, Trends in molecular medicine. 28, 194–209.

12. Murad, S., Michen, S., Becker, A., Fussel, M., Schackert, G., Tonn, T., Momburg, F. & Temme, A. (2022) NKG2C+ NK Cells for Immunotherapy of Glioblastoma Multiforme, International journal of molecular sciences. 23.

13. Breznik, B., Ko, M. W., Tse, C., Chen, P. C., Senjor, E., Majc, B., Habic, A., Angelillis, N., Novak, M., Zupunski, V., Mlakar, J., Nathanson, D. & Jewett, A. (2022) Infiltrating natural killer cells bind, lyse and increase chemotherapy efficacy in glioblastoma stem-like tumorospheres, Communications biology. 5, 436.

14. Du, Y., Pollok, K. E. & Shen, J. (2023) Unlocking Glioblastoma Secrets: Natural Killer Cell Therapy against Cancer Stem Cells, Cancers. 15.

15. Qiu, Z., Zhao, L., Shen, J. Z., Liang, Z., Wu, Q., Yang, K., Min, L., Gimple, R. C., Yang, Q., Bhargava, S., Jin, C., Kim, C., Hinz, D., Dixit, D., Bernatchez, J. A., Prager, B. C., Zhang, G., Dong, Z., Lv, D., Wang, X., Kim, L. J. Y., Zhu, Z., Jones, K. A., Zheng, Y., Siqueira-Neto, J. L., Chavez, L., Fu, X. D., Spruck, C. & Rich, J. N. (2022) Transcription Elongation Machinery Is a Druggable Dependency and Potentiates Immunotherapy in Glioblastoma Stem Cells, Cancer discovery. 12, 502–521.

16. Chava, S., Bugide, S., Gupta, R. & Wajapeyee, N. (2020) Measurement of Natural Killer Cell-Mediated Cytotoxicity and Migration in the Context of Hepatic Tumor Cells, Journal of visualized experiments : JoVE.

17. Bonanni, V., Antonangeli, F., Santoni, A. & Bernardini, G. (2019) Targeting of CXCR3 improves antimyeloma efficacy of adoptively transferred activated natural killer cells, Journal for immunotherapy of cancer. 7, 290.

18. Bernardini, G., Sciume, G., Bosisio, D., Morrone, S., Sozzani, S. & Santoni, A. (2008) CCL3 and CXCL12 regulate trafficking of mouse bone marrow NK cell subsets, Blood. 111, 3626–34.

19. Shaim, H., Shanley, M., Basar, R., Daher, M., Gumin, J., Zamler, D. B., Uprety, N., Wang, F., Huang, Y., Gabrusiewicz, K., Miao, Q., Dou, J., Alsuliman, A., Kerbauy, L. N., Acharya, S., Mohanty, V., Mendt, M., Li, S., Lu, J., Wei, J., Fowlkes, N. W., Gokdemir, E., Ensley, E. L., Kaplan, M., Kassab, C., Li, L., Ozcan, G., Banerjee, P. P., Shen, Y., Gilbert, A. L., Jones, C. M., Bdiwi, M., Nunez-Cortes, A. K., Liu, E., Yu, J., Imahashi, N., Muniz-Feliciano, L., Li, Y., Hu, J., Draetta, G., Marin, D., Yu, D., Mielke, S., Eyrich, M., Champlin, R. E., Chen, K., Lang, F. F., Shpall, E. J., Heimberger, A. B. & Rezvani, K. (2021) Targeting the alphav integrin/TGF-beta axis improves natural killer cell function against glioblastoma stem cells, The Journal of clinical investigation. 131.

